# Intrinsically aggregation-prone proteins form amyloid-like aggregates and contribute to tissue aging in *C. elegans*

**DOI:** 10.1101/417873

**Authors:** C. Huang, S. Wagner-Valladolid, A.D. Stephens, R. Jung, C. Poudel, T. Sinnige, M.C. Lechler, N. Schlörit, R.F. Laine, C.H. Michel, M. Vendruscolo, C.F. Kaminski, G.S. Kaminski Schierle, D.C. David

**Author notes:** These authors contributed equally to this work (S. Wagner-Valladolid, A.D. Stephens).

## Abstract

Reduced protein homeostasis and increased protein instability is a common feature of aging. Yet it remains unclear whether protein instability is a cause of aging. In neurodegenerative diseases and amyloidoses, specific proteins self-assemble into amyloid fibrils and accumulate as pathological solid aggregates in a variety of tissues. More recently, widespread protein aggregation has been described during normal aging, in the absence of disease processes. Until now, an extensive characterization of the nature of age-dependent protein aggregation and its consequences for aging has been lacking. Here, we show that age-dependent aggregates are rapidly formed by newly synthesized proteins and contain amyloid-like structures similar to disease-associated protein aggregates. Moreover, we demonstrate that age-dependent protein aggregation accelerates the functional decline of different tissues in *C. elegans*. Together, these finding reveal that the formation of amyloid aggregates is a generic problem of aging and likely to be an important target for strategies designed to maintain physiological functions in later stages of life.

## Introduction

Aging is a gradual decline in physiological functions and organ integrity. Diminished physical capacity and cognitive functions are already apparent before midlife, in the third decade of life (1). Significantly, the aging process greatly enhances the risk for chronic diseases leading to long-term disabilities in the elderly population and often premature death (2). A better understanding of what drives aging and in particular functional decline holds the promise of identifying targets to maintain quality of life with age. It is generally agreed that the ultimate root causes of aging occur at the molecular level (2). One of the most universal hallmarks of aging is increased protein instability (3, 4). In young individuals, an efficient protein homeostasis (proteostasis) network prevents the accumulation of protein instability. However, overwhelming evidence points to a collapse in the proteostasis network with age and consequently a decline in the ability to cope with protein instability (5). Still the role of protein instability in aging is unclear (6, 7).

During aging, an important cause of protein instability is cumulative damage through non-enzymatic posttranslational modifications occurring through reactions with metabolites and reactive oxygen species (4). Destabilizing mutations, transcriptional and translational inefficacy are also a significant source of protein instability (8, 9). In a disease context, a distinct form of protein instability namely protein aggregation is a common feature of amyloidoses and many neurodegenerative diseases. In these diseases, a specific protein converts from its native soluble conformation into an insoluble state by self-assembly into a cross-β sheet fibrillary structure termed amyloid. The amyloid fibrils accumulate as pathological deposits in a variety of different tissues. Although most proteins contain amyloid promoting sequences and will form amyloid fibrils in appropriate conditions *in vitro* (10, 11), there is little evidence for this conformation in metazoa in the absence of disease. Previously, we performed a global proteomic characterization of age-dependent protein insolubility, a hallmark of disease-associated protein aggregation, in wild-type *C. elegans* (12). We identified several hundred proteins enriched in distinct structural and functional characteristics that lose their native structure in aged animals and become highly-insoluble in strong detergents (12, 13). *In vivo* data demonstrate that these proteins accumulate in solid/immobile structures in aged animals. However, until now it is not known whether these age-dependent protein aggregates contain amyloid fibrils. Elucidating the structural nature of this novel type of protein instability would help to understand the causes and consequences of protein aggregation during aging and to explain potential interactions with disease-aggregating proteins.

Since the initial discovery in *C. elegans*, age-dependent protein aggregation has been demonstrated in different organs in mammals (14-18). Importantly, age-dependent protein aggregation is likely to be relevant in neurodegenerative processes. Indeed, the over-representation of proteins prone to aggregate with age among proteins sequestered in Alzheimer’s disease inclusions of amyloid plaques and tau neurofibrillary tangles highlights a possible contribution to pathological processes (12). Moreover, recent evidence shows that aggregates from wild-type aged mouse brains are a potent heterologous seed for amyloid-β (Aβ) aggregation (17). However, because of the difficulties to distinguish the effects of protein aggregation from other consequences of aging, it is still not clear whether age-dependent protein aggregation plays a role in accelerating the aging process (19). A previous study has shown that *C. elegans* treated with RNAi targeting genes encoding aggregation-prone proteins tend to live longer than randomly chosen targets (20). However, the interpretation of these results is complicated by the loss-of-function of these proteins. Recently, we found that aged animals with high aggregation levels of an RNA-binding protein (RBP) with a low-complexity prion-like domain were shorter lived, significantly smaller and less motile than animals with low RBP aggregation levels (13). Still this evidence does not provide a definite answer to whether age-dependent protein aggregation plays a causal role in aging rather than being a simple by-product. To understand whether protein aggregation is protective or harmful, it is informative to look at how long-lived animals modulate protein solubility. Longevity mechanisms and enhanced proteostasis are tightly coupled (5). However, whereas several studies show reduced age-dependent protein aggregation in long-lived animals (12, 13, 21), a recent study suggests that enhancing protein insolubility could be a strategy to promote longevity (22).

In the current study, we use fluorescence lifetime imaging (FLIM) (23) to reveal amyloid-like structures in age-dependent protein aggregates *in vivo*. Unlike protein instability caused by cumulative damage, age-dependent protein aggregates are formed by intrinsically aggregation-prone proteins shortly after their synthesis. Importantly, we demonstrate that age-dependent protein aggregation is toxic and contributes to functional decline in *C. elegans*.

## Results

### Age-dependent aggregating proteins are intrinsically prone to aggregate in certain tissues

Previously, we performed an extensive characterization of the aggregation of two proteins, casein kinase I isoform alpha (KIN-19) and Ras-like GTP-binding protein rhoA (RHO-1). Both KIN-19 and RHO-1 were identified among the proteins with the highest propensity to become insoluble with age in wild-type *C. elegans* somatic tissues (12). *In vivo* analysis of animals expressing these proteins fused to fluorescent tags showed the appearance of highly immobile deposits with age (12). Among the insoluble proteome, the enrichment of certain structural features such as high aliphatic amino acid content or propensity to form β-sheet-rich structures shows that age-dependent protein aggregation is not random (12, 13). To understand whether KIN-19 and RHO-1 have an intrinsic capacity to aggregate similar to disease-associated proteins or whether a progressive accumulation of protein damage caused by non-enzymatic posttranslational modifications is required to induce their aggregation, we evaluated the dynamics of protein aggregation *in vivo*. Protein labeling with mEOS2, a green-to-red photoconvertible fluorescent protein, has been successfully used to track protein dynamics (24). In the present case, we used the mEOS2 tag to investigate how fast newly synthesized KIN-19 and RHO-1 aggregate. For this, we generated transgenic animals expressing KIN-19∷mEOS2 in either the pharynx or in the body-wall muscles and transgenic animals expressing RHO-1∷mEOS2 in the pharynx. The mEOS2 tag did not disrupt the aggregation potential of KIN-19, as the absence of fluorescence recovery after photobleaching confirms that both KIN-19∷mEOS2 puncta in the pharynx and body-wall muscle are highly immobile structures (Figure S1 A,B).

To follow newly synthesized proteins, we set-up a system to perform irreversible photoconversion of the mEOS2 tag present in live animals from green to red by exposing them to intense blue fluorescent light. At a defined time-point, we photoconverted the mEOS present in aggregates to red. After the photoconversion, newly synthesized proteins emitted green fluorescence and could thus easily be distinguished from old (photoconverted/red) aggregates. This method allowed us to follow the rate of new aggregate formation and the rate of old aggregate removal in a population of transgenic animals over time. Of note, we observed photoconversion of all aggregates, however the core region of some larger aggregates continued to emit green fluorescence (Figure 1 A, B). Interestingly, we observed a doubling in the number of animals with newly-formed green aggregates, 24 hours after the conversion at day 5 of adulthood in both the pharyngeal and body-wall muscles. Conversely, analysis of the red aggregates suggested a slow removal of old aggregates (Figure 1 A, B). Confocal imaging shows that newly synthesized KIN-19 associate with pre-existing aggregates (Figure 1 C). However, we also observed large aggregates emitting only green fluorescence suggesting that seeding is not a prerequisite for KIN-19 aggregate formation (Figure 1 C). We observed similar aggregation kinetics in animals with RHO-1∷mEOS2 expressed in the pharynx (Figure S1 C). Together, these results reveal that aggregate formation proceeds rapidly after protein synthesis.

**Figure 1:**
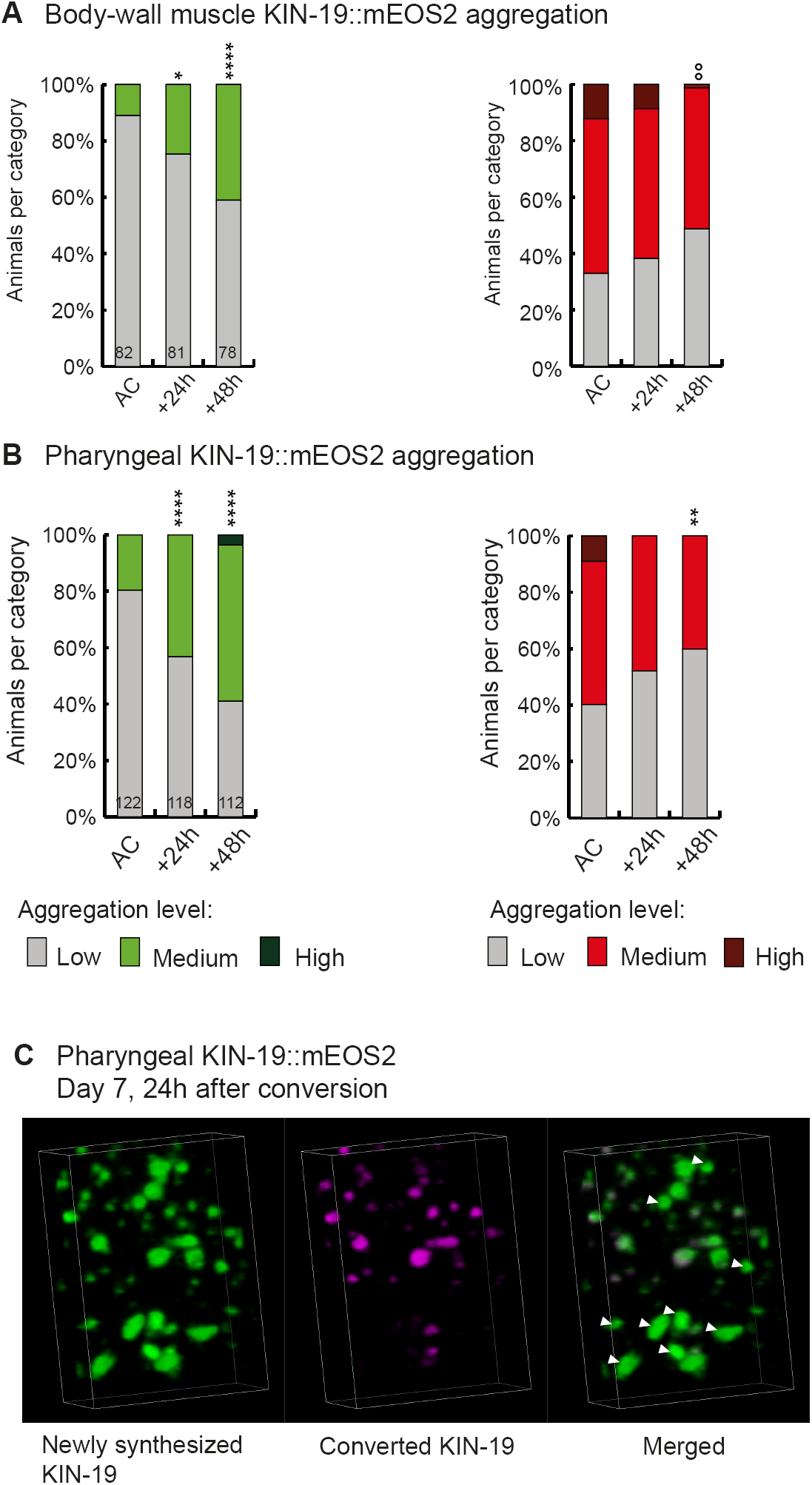
Newly synthesized KIN-19 rapidly transitions into aggregates in aged animals. (A, B) Following photoconversion at day 5 in the pharynx or in the body-wall muscle, the number of animals with newly synthesized green-emitting non-converted KIN-19∷mEOS2 aggregates doubles over 24 hours. Conversely levels of red-emitting converted aggregates slowly declines. Aggregation evaluated in *Pkin-19∷KIN-19∷mEOS2* and *Pmyo-3∷KIN-19∷mEOS2* transgenic animals. Numbers of worms indicated in the bars. Fisher’s exact test comparing low versus medium + high aggregation levels to after conversion (AC): *p < 0.05, **p < 0.01, ****p < 0.0001. Fisher’s exact test comparing low + medium versus high aggregation levels to after conversion (AC): °°p < 0.01. (C) 24 hours after photoconversion at day 7, newly synthesized KIN-19∷mEOS2 (green emitting) forms new aggregates and associates around older aggregates (red emitting). 3D reconstruction of the pharyngeal anterior bulb region. Arrow heads highlight large new aggregates formed independently of previous aggregates.

### Age-dependent protein aggregates contain amyloid-like structures *in vivo*

Both KIN-19 and RHO-1 proteins are nearly identical to their human orthologues (sequence identity over 87%). X-ray diffraction shows that the human orthologues are globular proteins formed by a series of α-helices and β-strands. The ability of KIN-19 and RHO-1 to aggregate shortly after synthesis suggests that their aggregation may be driven by folding intermediates. Notably, both proteins have several segments with a high amyloid propensity (Figure S2 A, B) (10). This raises the possibility that KIN-19 and RHO-1 gain an amyloid-like conformation during their aggregation with age. We have previously shown that FLIM can be used to determine whether certain proteins are likely to form amyloid-like aggregates as the formation of amyloid fibrils leads to significant drop in the fluorescence lifetime of the fluorescent-tagged amyloid proteins due to quenching (23, 25, 26). In the current study, we applied FLIM to determine whether KIN-19 and RHO-1 form amyloid-like aggregates in live *C. elegans*. For this, we generated transgenic animals expressing translational fusions with the yellow fluorescent protein Venus in the pharynx. We first confirmed that there is an age-dependent increase in aggregate formation by KIN-19∷Venus and RHO-1∷Venus in the pharynx (Figure S3 A, B) (13). Using FLIM, we found that pharyngeal KIN-19 and RHO-1∷Venus display a significantly decreased fluorescence lifetime compared to Venus only control worms (Figure 2 A-C). Areas with the strongest drop in fluorescence lifetime co-localize with fluorescent-labeled aggregates (Figure 2 D). In both models, we observed a significant drop in the fluorescence lifetime already at day 1, consistent with the appearance of aggregates in young animals due to protein overexpression. In particular, between day 1 and day 7 of adulthood, we measured a dramatic decrease in fluorescence lifetime of RHO-1 aggregates and a more modest decrease in the fluorescence lifetime of KIN-19 aggregates. This is consistent with the age-dependent increase in aggregate formation by both proteins and the fact that RHO-1 puncta tend to be larger and more solid than KIN-19 puncta. Indeed, all RHO-1 puncta evaluated showed no recovery after photobleaching whereas 30% of KIN-19 puncta showed some recovery as previously described (Figure S3 C, D compared to (12)).

**Figure 2:**
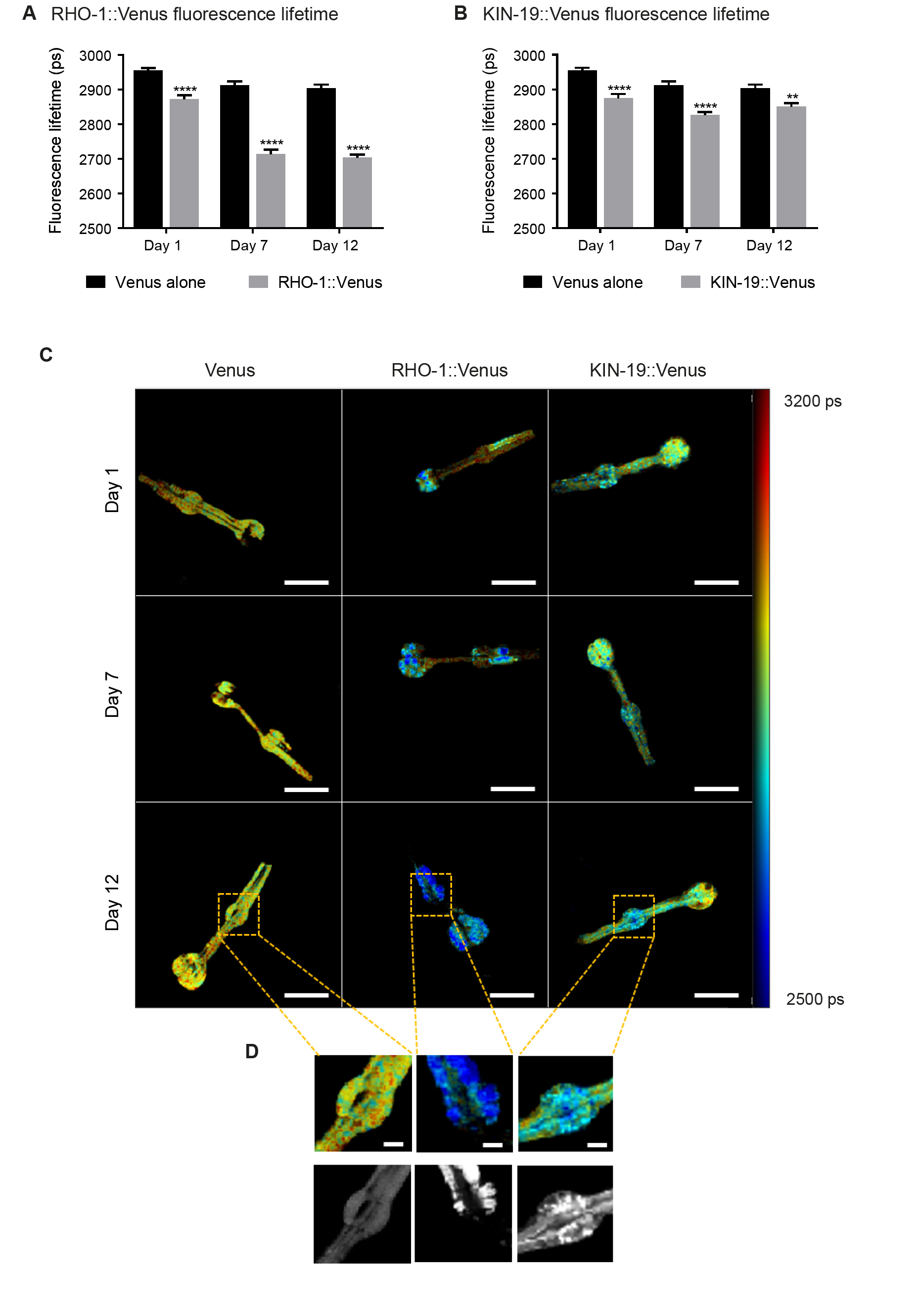
Drop in fluorescence lifetime reveals amyloid-like structure of KIN-19 and RHO-1 aggregates in live *C. elegans*. (A) Bar diagram showing the intensity-averaged mean fluorescence lifetime values of day 1 (2872 ± 11 ps), day 7 (2713 ± 13 ps) and day 12 (2704 ± 8 ps) adult worms, expressing RHO-1∷Venus in the pharynx compared to worms expressing Venus only. n = 7-10, 2 repeats. (B) Bar diagram showing the intensity-averaged mean fluorescence lifetime images of day 1 (2875 ± 12 ps), day 7 (2826 ± 9 ps) and day 12 (2851 ± 10 ps) adult worms, expressing KIN-19∷Venus in the pharynx compared to worms expressing Venus only. n = 7-10, 2 repeats. (C) Representative intensity-averaged fluorescence lifetime images for Venus only, RHO-1∷Venus and KIN-19∷Venus worms. Scale = 50 μm. (D) Zoom into anterior bulb for Venus only, RHO-1∷Venus and KIN-19∷Venus worms. Scale = 10 μm. Data is shown as mean lifetime + SEM and the statistical analysis was performed using two-way ANOVA with Sidak’s multiple comparisons test ** = p<0.01, **** = p<0.0001.

To gain more insight into the capacity of RHO-1 to form amyloid fibrils, we expressed and purified recombinant RHO-1 (Figure S4 A, B) and aggregated it over time by constant shaking in stabilizing buffer to induce fibrillisation at 37°C for a week. We obtained low quantities of RHO-1 fibrils, which we were able to analyze by transmission electron microscopy (TEM). Analysis of RHO-1 by TEM revealed fibril-like structures which resemble amyloid fibrils such as formed by Huntingtin (Htt) (Figure S4 C) (27). Indeed, RHO-1 fibrils had a morphology more similar to Htt fibrils which are shorter and more branch-like compared to fibrils formed from other amyloids such as alpha-synuclein which are long and flexible. Additionally, RHO-1 and Htt fibrils display no twisting which can be observed in alpha-synuclein fibrils, suggesting that the packing of the monomeric structure and the interaction between protofibrils may be different (28). Importantly, already at day 1, we found similar fibrillary structures in extracts from RHO-1∷Venus transgenic animals (Figure S4 D compared to S4 E).

Together, these findings strongly suggest that Casein kinase I isoform alpha and Ras-like GTP-binding protein rhoA aggregates contain amyloid-like structures *in vivo*.

### Age-dependent protein aggregation accelerates functional decline

A reliable indicator of human aging is a decline in physical capacity such as decreased muscle strength and coordination (1). Similarly, *C. elegans* displays an age-related decline in pharyngeal pumping and body movement (29). To investigate whether age-dependent protein aggregation accelerates functional decline with age, we measured pharyngeal pumping and swimming in liquid (thrashing) in transgenic animals overexpressing fluorescent-labeled KIN-19 and RHO-1 either in the pharynx or the body-wall muscles. In all models, protein aggregation increased with age, as measured by the change from a diffuse distribution to the formation of specific puncta by the fluorescent-tagged proteins (Figure S5). Among the transgenic models generated, the largest age-dependent changes were observed for *C. elegans* expressing KIN-19∷tagRFP in the pharynx (Figure S5 A, E) (12). Whereas the majority of these transgenic animals have no aggregation at day 2, this dramatically changes with age and at day 8, the majority display high levels of KIN-19∷tagRFP aggregation. Importantly, pumping frequency was strongly reduced in day 7 aged animals with the highest levels of pharyngeal KIN-19∷tagRFP aggregation compared to those with the lowest aggregation levels (Figure 3 A, Table S1). The detrimental effect of age-dependent protein aggregation was also apparent in animals with KIN-19∷tagRFP aggregation in the body-wall muscles, since we observed an earlier decline in swimming frequency associated with KIN-19∷tagRFP aggregation in the body-wall muscle (Figure S6 A). This effect was amplified in animals with the highest level of aggregation (Figure 3 B). Of note, the fluorescent tagRFP expressed alone in the pharynx or in the body-wall muscle did not affect muscle function (Figure S6 A, C).

**Figure 3:**
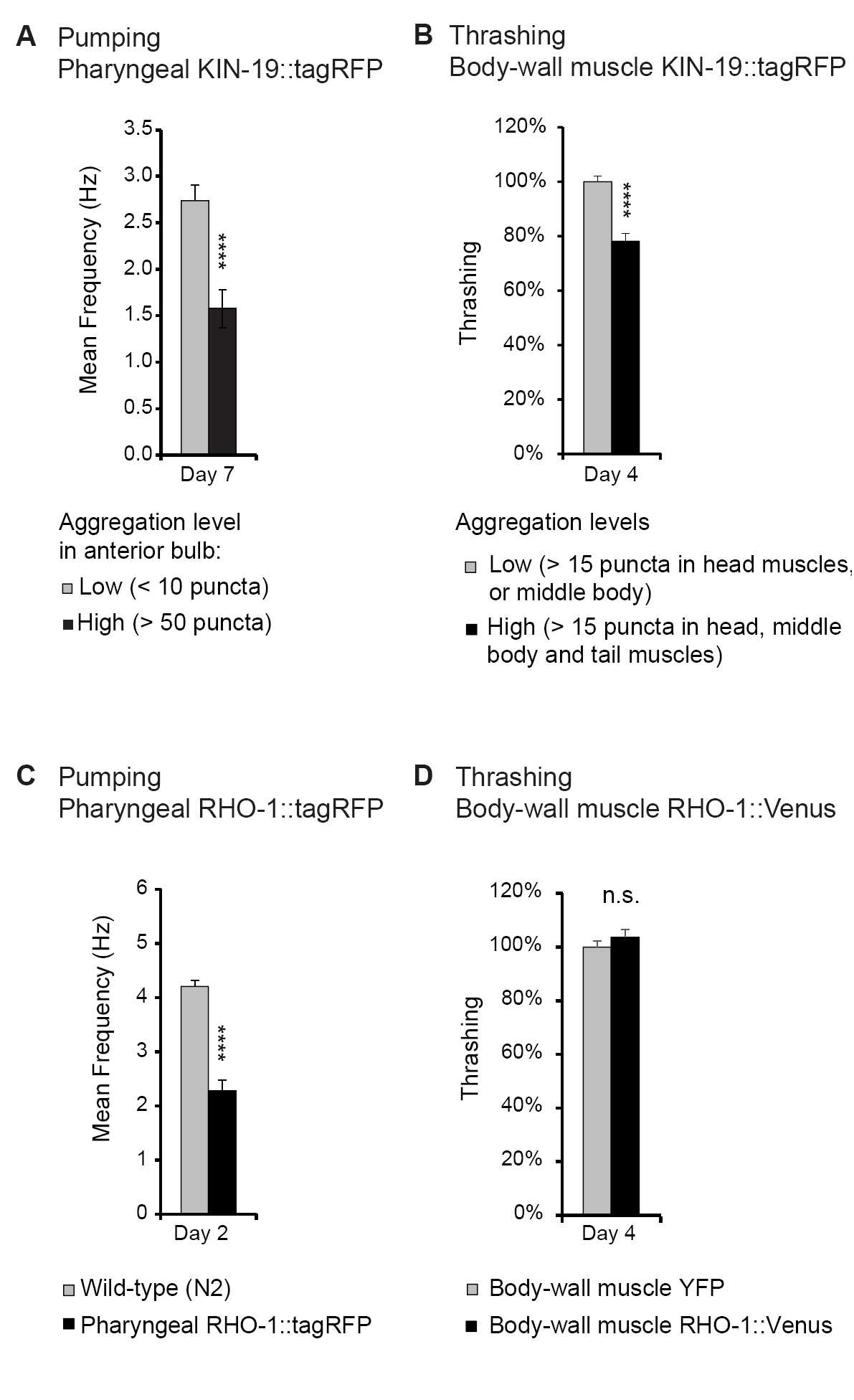
Age-dependent protein aggregation impairs pharyngeal and body-wall muscle function. (A) Aged animals with high levels of pharyngeal KIN-19 aggregation have reduced pharyngeal pumping. N= 21-28 animals analyzed per group. T-test: p< 0.0001. (B) Aged animals with high levels of KIN-19 aggregation in the body-wall muscle display reduced thrashing. Mean body bends per seconds are set to 100% in animals with low aggregation. Mann-Whitney test: p< 0.0001. (C) Young animals with RHO-1 aggregation in pharynx have impaired pharyngeal pumping. N= 25-27 animals analyzed per group. T-test: p< 0.0001. (D) Overexpression of RHO-1 without aggregation in the body-wall muscles does not influence thrashing. Mean body bends per seconds are set to 100% in *Punc-54∷yfp* transgenic animals. Mann-Whitney test: non-significant. SEM represented, data in Table S1.

To further show that protein aggregation is the cause of the functional decline in our transgenic animal models rather than other co-occurring aging factors, we examined the effects of protein aggregation in the absence of aging. Most likely due to the high level of overexpression in the pharyngeal muscles, RHO-1∷tagRFP aggregated abundantly already in young animals (Figure S5 C, E). We found that these high levels of RHO-1∷tagRFP aggregation strongly impaired pharyngeal pumping in young animals (day 2) (Figure 3 C). Notably, these young animals displayed pumping rates normally observed only in aged animals. To exclude that RHO-1 overexpression itself is toxic to *C. elegans*, we examined another transgenic model where RHO-1∷Venus is expressed under the strong *C. elegans* body-wall muscle promoter *Punc-54*. These animals have similar levels of transgene expression compared to animals with body-wall muscle KIN-19∷tagRFP (Figure S5 F). Yet, whereas KIN-19 aggregates abundantly in the body-wall muscle, RHO-1∷Venus hardly aggregates in this tissue (Figure S5 D). Thus, these transgenics are a suitable control to identify RHO-1 toxicity caused by overexpression. Importantly, we did not observe reduced thrashing in animals with body-wall muscle RHO-1, demonstrating that overexpressed of RHO-1 without concomittant aggregation is not toxic (Figure 3 D). Similarly, the observation that only animals with the highest levels of pharyngeal KIN-19∷tagRFP aggregation have functional impairment shows that KIN-19∷tagRFP overexpression alone does not cause pumping defects (Figure 3 A, S6 B). Together, these positive and negative controls reveal that protein aggregation itself is detrimental in *C. elegans*.

Collectively, these findings show that animals with accelerated protein aggregation experience an earlier onset of functional decline in the tissues affected.

## Discussion

Widespread protein aggregation in the context of normal aging has been observed in *C. elegans* (12, 20, 22), *Drosophila* (21), *Saccharomyces cerevisiae* (30) and in mammals, notably in neural stem cells (16), heart (15) and skeletal muscles (18), bone marrow and spleen (14). Whereas these hundreds of proteins are maintained in a functional and soluble state in young animals, they lose their functional structure with age and accumulate into highly-insoluble aggregates. The detergent insoluble properties and solid nature of the aggregates indicate similarities with disease-associated protein aggregation. However, until now it has not been known whether age-dependent protein aggregates display amyloid-like structures, a key characteristic of disease-associated protein aggregation. In this study, we focus on two normally globular proteins, Casein kinase I isoform alpha and Ras-like GTP-binding protein rhoA. We show that aggregates of either proteins display distinct fluorescence quenching properties characteristic of amyloid structures. In neurodegenerative diseases and amyloidosis, protein aggregation is a crucial part of the pathological process. We demonstrate that aggregates formed by proteins prone to insolubility with age lead to accelerated functional decline in the tissues affected. Our results predict that even if only a proportion of the hundreds of proteins becoming insoluble with age form harmful amyloid-like aggregates, this would be a significant cause of tissue aging for a variety of organs.

The speed of aggregate formation and the presence of amyloid-like structure brings significant insight into the aggregation process occurring during normal aging. We demonstrate that newly synthesized proteins rapidly assemble into large aggregates. The recruitment of newly synthesized proteins into existing aggregates indicates a seeding effect. However, we also observed the rapid formation of new aggregates entirely made up of newly synthesized proteins. This shows that molecular aging of the protein caused by progressive accumulation of damage is not required for proteins to aggregate with age. Moreover, amyloid-like aggregates forming already in young animals indicate an intrinsic aggregation capacity. Therefore, it is likely that these proteins self-assemble into amyloid structures directly from the unfolded state occurring during or shortly after translation rather than undergoing unfolding from their natively folded state prior to amyloid formation. This is consistent with protein folding intermediates being particularly at risk of aggregating as exemplified by a recent study revealing that newly synthesized proteins constitute the majority of the insoluble fraction prompted by thermal stress (31). Furthermore, artificially increasing ribosome pausing during translation causes widespread protein aggregation (9). Both Casein kinase I isoform alpha and Ras-like GTP-binding protein rhoA contain several hexapeptides with high propensity for fibrillation which are normally buried in the fully folded protein (Figure S2). Interestingly, the most prominent amyloid-promoting sequence for both proteins is localized near the N-terminus. Therefore, aberrant interactions may start already during translation. As protein aggregation typically results in loss of function, there is a strong evolutionary pressure to avoid this and chaperones have evolved to recognize specifically sequences of high aggregation propensity (32). Our findings predict that age-dependent protein aggregation would result from decreased levels of chaperones linked to protein synthesis rather than chaperones induced by stress (33, 34). Impaired proteasome-mediated removal of unfolded proteins directly after synthesis (35) could also significantly contribute to age-dependent protein aggregation. Interestingly, the relative lack of Ras-like GTP-binding protein rhoA aggregation in the body-wall muscles compared to the pharyngeal muscles shows that this protein only aggregates within a specific cellular environment. Thus tissue-specific factors could be crucial for the aggregation process. These could be differences in the tissue proteostasis network and reliance on certain proteostasis components with age (36-38) but also age-related changes in the local tissue environment. For example, tissue-specific changes in ATP levels (39), availability of certain ions such as Ca^2+^ (40), redox state (41), could explain why certain proteins aggregate in one tissue but not another.

The resemblance between age-dependent protein aggregation and disease-associated protein aggregation raises the question whether they have similar proteotoxic mechanisms. Our results show that there is a strong correlation between the presence of large aggregates and impaired tissue function. In disease, intracellular aggregate toxicity is caused, at least in part, by sequestration and the resulting loss of function of essential cellular proteins (42-44). Notably, proteostasis network components are significantly enriched in the age-dependent insoluble proteome (12). Therefore loss-of-function of proteins responsible for preventing aggregation could be a key source of toxicity. The correlation between amyloid-like aggregates and proteotoxicity observed in the present study, does not exclude that intermediate forms on the path to age-dependent protein aggregation are also proteotoxic. Indeed, disease-associated pre-fibrillar species or oligomers can be highly toxic by interacting through their hydrophobic side chains with other cellular components and in particular lipid membranes (45, 46). Further experiments will be needed to characterize which types of intermediate species occur during age-dependent protein aggregation and to evaluate their potential toxicity. Conversely, it remains possible that some forms of age-dependent protein aggregation could be protective or manipulated to form harmless non-amyloid aggregates. Finally, it is possible that age-dependent protein aggregation occurring in one tissue will induce accelerated aging in another tissue. Following the evidence for seeding, self-propagation and cell-to-cell transfer of amyloid aggregating species in a disease context (47), it is intriguing to speculate that the same mechanisms may arise with proteins aggregating during normal aging.

The proteotoxicity and amyloid conformation of age-dependent protein aggregation has important implications for diseases associated with protein aggregation. Accelerated functional decline and the overload of the proteostasis network caused by age-dependent protein aggregation could indirectly enhance disease-associated pathogenesis. There is also evidence for a direct connection between disease and aging-related aggregation. Indeed, a significant proportion of the proteins sequestered in disease pathological deposits are prone to aggregate with age (12, 48). Notably, Casein kinase I isoform alpha is present in tau aggregates (49, 50). Therefore, age-dependent aggregation-prone proteins interact with disease-aggregating proteins in humans. Furthermore, we have demonstrated that minute amounts of insoluble proteins from aged wild-type mouse brains or aged *C. elegans* is sufficient to cross-seed amyloid-β aggregation *in vitro* (17). One possibility is that the highly hydrophobic nature of the amyloid-like structures in age-dependent protein aggregates provides a destabilizing surface that can promote the conformational conversion of disease-associated aggregating proteins. If the composition of the insoluble proteome is cell-specific, increases in age-dependent protein aggregation, for example through higher expression or somatic aggregation-promoting mutations (51, 52), could sensitize specific cells to disease-associated pathogenesis.

In summary, our study demonstrates that aggregation of proteins during normal aging strongly resembles pathological protein aggregation observed in neurodegenerative diseases and amyloidosis. We show that age-dependent protein aggregation causes early functional decline, a read-out of accelerated aging. These findings emphasize age-dependent protein aggregation as an important target to restore physical capacity and to promote healthy aging. Already promising results reveal that lysosome activation in the germline and in aged neural stem cells clears protein aggregates and rejuvenates the cells (16, 53).

## Methods

### Transgenics

CF3166: *muEx473[Pkin-19∷kin-19∷tagrfp + Ptph-1∷gfp]*

CF3649: N2; *muIs209[Pmyo-3∷kin-19∷tagrfp + Ptph-1∷gfp]*

CF3706: N2; *muEx587[Pkin-19∷kin-19∷meos2 + Punc-122∷gfp]*

DCD13: N2; *uqIs9[Pmyo-2∷rho-1∷tagrfp + Ptph-1∷gfp]*

DCD69: N2; *uqEx4[Pmyo-3∷kin-19∷meos2]*

DCD83: *ttTi5605II; unc-119(ed3)III; uqEx[Pmyo-2∷rho-1∷meos2 + Punc-122∷gfp + cb-unc-119(+)]*

DCD127: N2; *uqEx22[Punc-54∷rho-1∷venus]*

DCD146: N2; *uqIs12[Pmyo-2∷rho-1∷venus]*

DCD179: N2; *uqEx37[Pkin-19∷kin-19∷venus + punc-122∷gfp]*

AM134: *rmIs126[Punc-54∷YFP]*

CL2122: *dvIs15 [(pPD30.38) unc-54(vector) + (pCL26) mtl-2∷GFP].*

### Maintenance

All strains were kept at 15°C on NGM plates inoculated with OP50 using standard techniques. Age-synchronization was achieved by transferring adults of the desired strain to 20°C and selecting their progeny at L4 stage. All experiments were performed at 20°C. Day 1 of adulthood starts 24h after L4.

### Imaging

For confocal analysis using a Leica SP8 confocal microscope with the HC PL APO CS2 63×1.40 oil objective, worms were mounted onto slides with 2% agarose pads using 2 μM levamisole for anaesthesia. Worms were examined using the Leica HyD hybrid detector. The tag mEOS2 was detected using 506nm as excitation and an emission range from 508-525nm for green fluorescence and 571nm as excitation and an emission range from 573-602nm for red fluorescence. 3D reconstructions were performed using the Leica Application Suite (LAS X). FRAP analysis was performed as previously described (12) using the Leica SP8 confocal microscope with the HC PL APO CS2 63x 1.30 glycerol objective and PMT detector.

### Photoconversion of mEOS2-tag and quantification of fluorescence levels

For photoconversion, worms were transferred onto a small (diameter 35 mm) NGM plate without food. The plate was placed 0.5 cm below a collimator (Collimator High-End Lumencor, Leica, Germany) fitted with a filter for blue fluorescence (387/11 BrightLine HC, diameter 40mm) and illuminated by a Lumencor Sola SE II (AHF, Tübingen). Conversion of mEOS2 in transgenic animals was performed four times for five minutes, with 2 minutes pauses between exposures.

### Aggregation quantification *in vivo*

Aggregation levels were determined using Leica fluorescence microscope M165 FC with a Planapo 2.0x objective. Animals expressing *Pkin-19∷KIN-19∷mEOS2, Pkin-19∷KIN-19∷Venus or Pkin-19∷KIN-19∷TagRFP* were divided into less than 10 puncta (low aggregation), between 10 and 100 puncta (medium aggregation) and over 100 puncta in the anterior bulb (high aggregation). Animals overexpressing *Pmyo-2∷ RHO-1∷Venus* were divided into less than 10 puncta in anterior or posterior pharyngeal bulb (low aggregation), over 10 puncta in either bulbs (medium aggregation) and over 10 puncta in both bulbs (high aggregation). Because of extensive RHO-1 aggregation in animals overexpressing *Pmyo-2∷RHO-1∷TagRFP*, aggregation was only quantified in the isthmus: animals with no aggregation (low aggregation), animals with aggregation in up to 50% (medium aggregation) and animals with aggregation in more than 50% (high aggregation) of the isthmus. Animals overexpressing *Pmyo-3∷KIN-19∷TagRFP* were divided into three categories: animals with more than 15 puncta in the head or the middle body region (low aggregation), animals with more than 15 puncta in the head and the middle body region (medium aggregation) and animals with puncta in head, middle body and tail region (high aggregation). Counting was done in a blind fashion. Two-tailed Fisher’s exact test using an online tool (https://www.graphpad.com/quickcalcs/contingency1.cfm) was performed for statistical analysis.

### Fluorescence lifetime imaging *in vivo*

For fluorescence lifetime imaging, transgenic *C. elegans* were mounted on microscope slides with 2.5% agarose pads using 25 mM NaN3 as anaesthetic. All samples were assayed on a modified confocal-based platform (Olympus FV300-IX70) equipped with a 60x oil objective (PLAPON 60XOSC2 1.4NA, Olympus, Germany) and integrated with time-correlated single photon counting (TCSPC) FLIM implementation. A pulsed supercontinuum (WL-SC-400-15, Fianium Ltd., UK) at 40MHz repetition rate served as the excitation source. YFP was excited at 510nm using a tuneable filter (AOTFnC-400.650, Quanta Tech, New York, USA). The excitation light was filtered with FF03-510/20 and the fluorescence emission was filtered with FF01-542/27 (both bandpass filters from Semrock Inc., New York, USA) before reaching the photomultiplier tube (PMC-100, Becker & Hickl GmBH, Berlin, Germany). Photons were recorded by a SPC-830 (Becker and Hickl GmBH, Germany) module that permits sorting photons from each pixel into a histogram according to the photon arrival times. Photons were acquired for two minutes to make a single 256 X 256 FLIM image and photobleaching was verified to be negligible during this time. Photon count rates were always kept below 1% of the laser repetition rate to avoid pulse pileup. All raw FLIM images were fitted with a single exponential decay function using FLIMfit (54) and exported to MATLAB (Mathworks, Inc., Natick, Massachusetts, USA) to obtain an intensity weighted lifetime average for each image. Statistical analysis was carried out using two-way ANOVA followed by Sidak’s multiple comparisons test in Graphpad Prism software (La Jolla, California, USA).

### Pharyngeal pumping analysis

Electrical activity of the pharyngeal pumping was measured using the NemaMetrix ScreenChip System (NemaMetrix, Eugene OR). The recordings were analysed by NemAnalysis v0.2 software and student’s t test was used for statistical analysis. Additional details are included in the supplementary methods.

### Thrashing analysis

To quantify movement in terms of body-bends-per-second, movies of worms swimming in liquid were acquired with high frame rates (15 frames per second) using a high-resolution monochrome camera (JAI BM-500 GE, Stemmer imaging GmbH, Puchheim, Germany). Five consecutive 30-second movies were made for each group of worms. The movies were then analysed using the ImageJ wrMTrck plugin (55). Mann-Whitney test was used for statistical analysis (GraphPad Prism 7).

Analysis of *in vitro* RHO-1 fibrils and *in vivo* RHO-1 fibrils from worm lysates:

Purification of recombinant RHO-1 and fibrillisation, extraction of RHO-1 fibrils from worm lysates, analysis by transmission electron microscopy are described in the supplementary methods.

## Acknowledgements

This work was supported by funding from the DZNE and a Marie Curie International Reintegration Grant (322120 to D.C.D). K.C.F. acknowledges funding from the UK Engineering and Physical Sciences Research Council (EPSRC). G.S.K.S. and K.C.F. acknowledge funding from the Wellcome Trust, the UK Medical Research Council (MRC), Alzheimer Research UK (ARUK), and Infinitus China Ltd. S.A.D. acknowledges Alzheimer Research UK (ARUK) travel grants. Some strains were provided by the CGC, which is funded by NIH Office of Research Infrastructure Programs (P40 OD010440).

## Author contributions

C.H., D.C.D., G.S.K.S. designed experiments; C.H., W.V.S., S.A.D., R.J., M.C.L, S.T., L.R.F., M.C.H. performed research; C.H., W.V.S., S.A.D., R.J., P.C. analysed data; G.H., P.C., M.V., K.C.F. contributed new reagents/analytic tools; C.H., W.V.S., S.A.D., S.T., G.S.K.S., D.C.D wrote the paper.

## Figure legends

**Supplementary figure 1:**
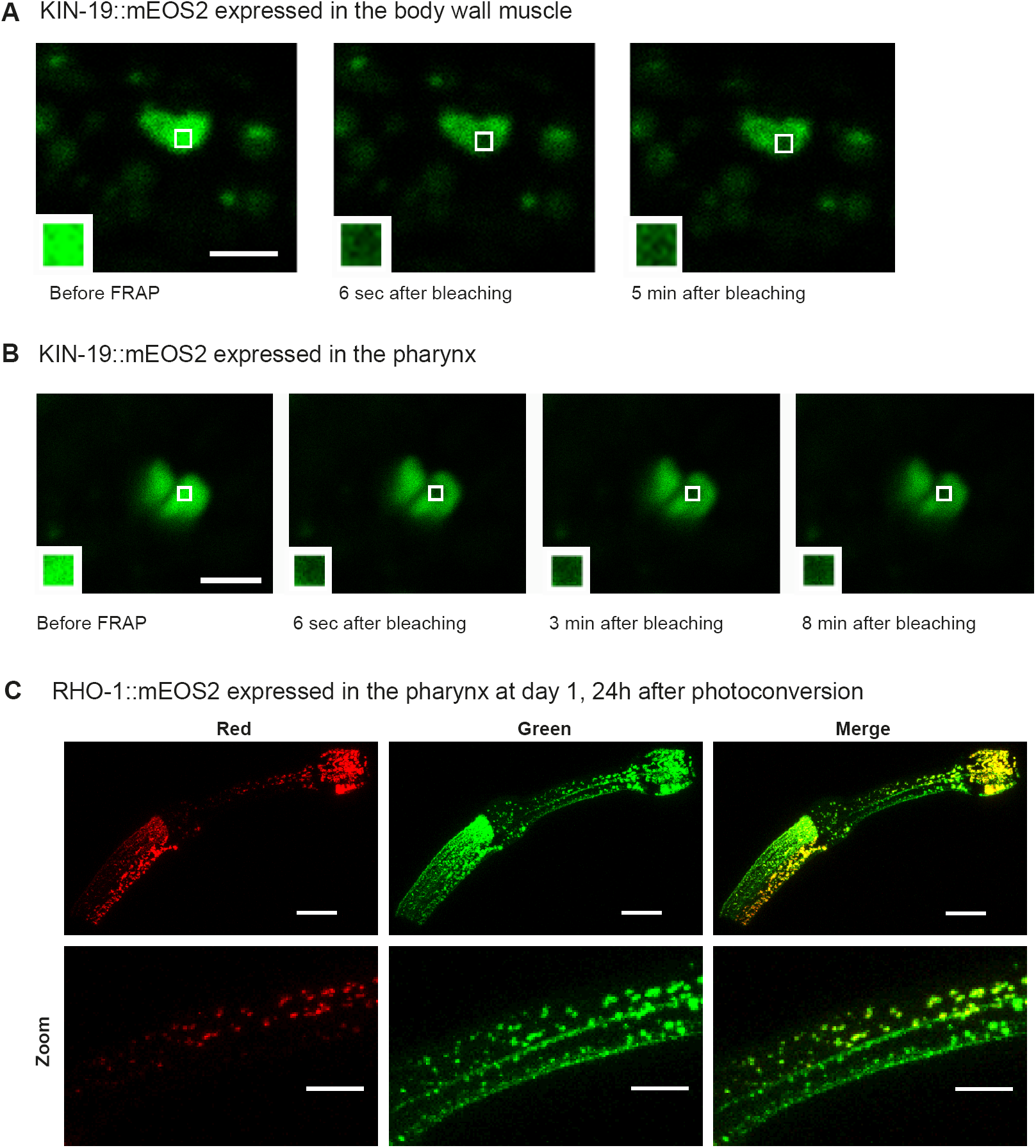
Newly synthesized RHO-1 rapidly transitions into aggregates in young animals. (A) KIN-19∷mEOS2 puncta visible in the body wall muscle contain immobile protein, demonstrated by lack of fluorescent recovery after photobleaching (FRAP) after 5 minutes. Area bleached enlarged in caption. Scale bar 2 μm. (B) KIN-19∷mEOS2 puncta visible in the pharynx contain immobile protein, demonstrated by FRAP after 8 minutes. Area bleached enlarged in caption. Scale bar 2 μm. (C) RHO-1∷mEOS2 aggregates strongly already at day 1 and forms new aggregates (green) 24 h after photoconversion and associates with older aggregates (red). Scale bar 15 μm, 7 μm in zoom.

**Supplementary figure 2:**
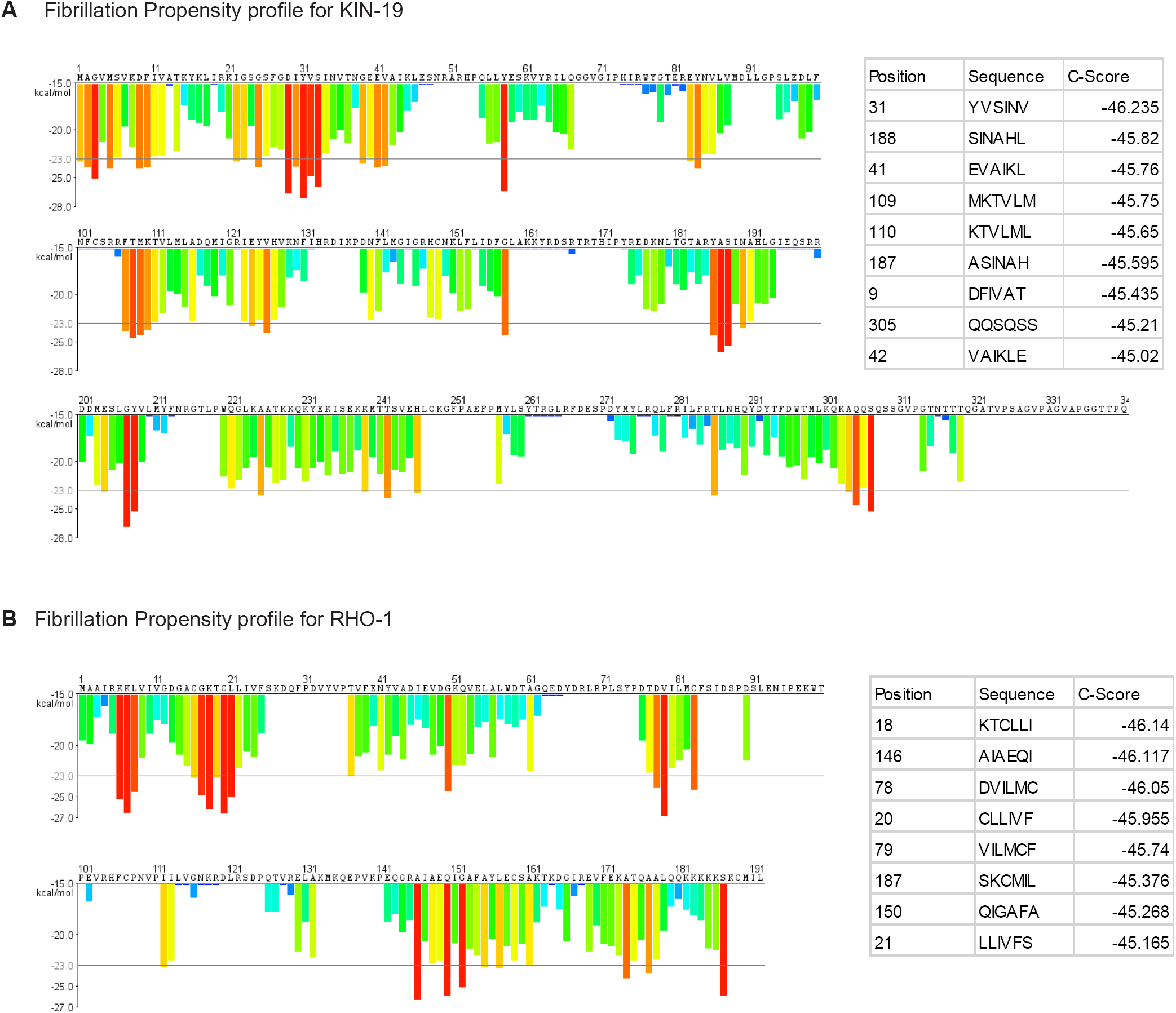
Fibrillation propensity calculate by ZipperDB. (A) Fibrillation propensity profile for KIN-19 (B) Fibrillation propensity profile for RHO-1 Table inlet shows hexapeptides with Composite scores below -45. ZipperDB: https://services.mbi.ucla.edu/zipperdb/

**Supplementary figure 3:**
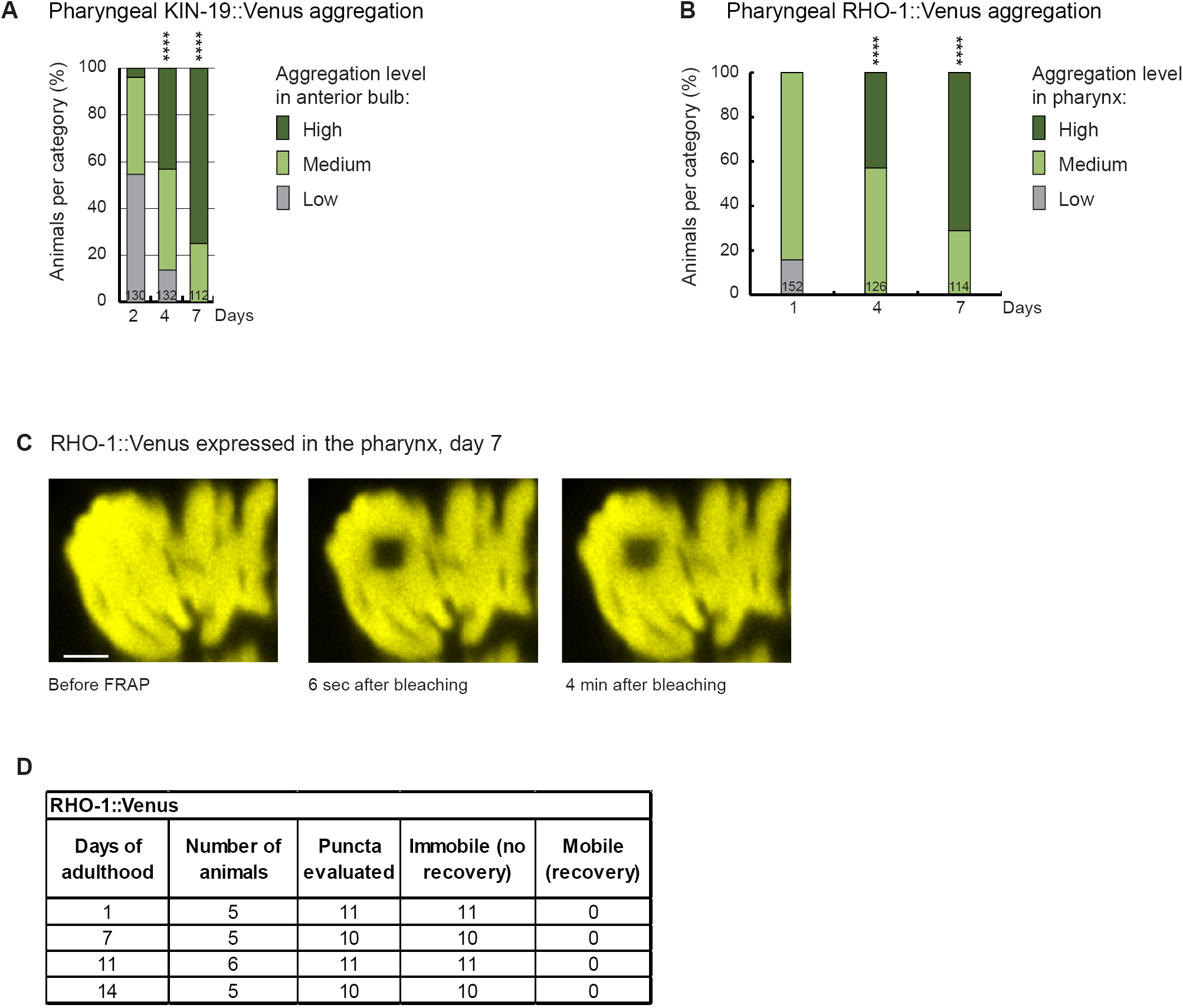
Aggregation with age of RHO-1∷Venus and KIN-19∷Venus. (A) Increased pharyngeal KIN-19 aggregation with age in animals expressing *Pkin-19∷KIN-19∷Venus*. Panel reproduced from https://doi.org/10.1016/j.celrep.2016.12.033. (B) Increased pharyngeal RHO-1 aggregation with age in animals expressing *Pmyo-2∷RHO-1∷Venus*. Numbers of worms indicated in the bars. Significance calculated low + medium versus high aggregation levels compared to day 2 of adulthood; Fisher’s exact test: ****p< 0.0001 (C) RHO-1∷Venus puncta visible in the pharynx contain immobile protein, demonstrated by absence of fluorescence recovery after 4 minutes. Scale bar 2 μm. (D) RHO-1∷Venus aggregates are highly immobile. Table summarizes FRAP experiments carried out at different ages.

**Supplementary figure 4:**
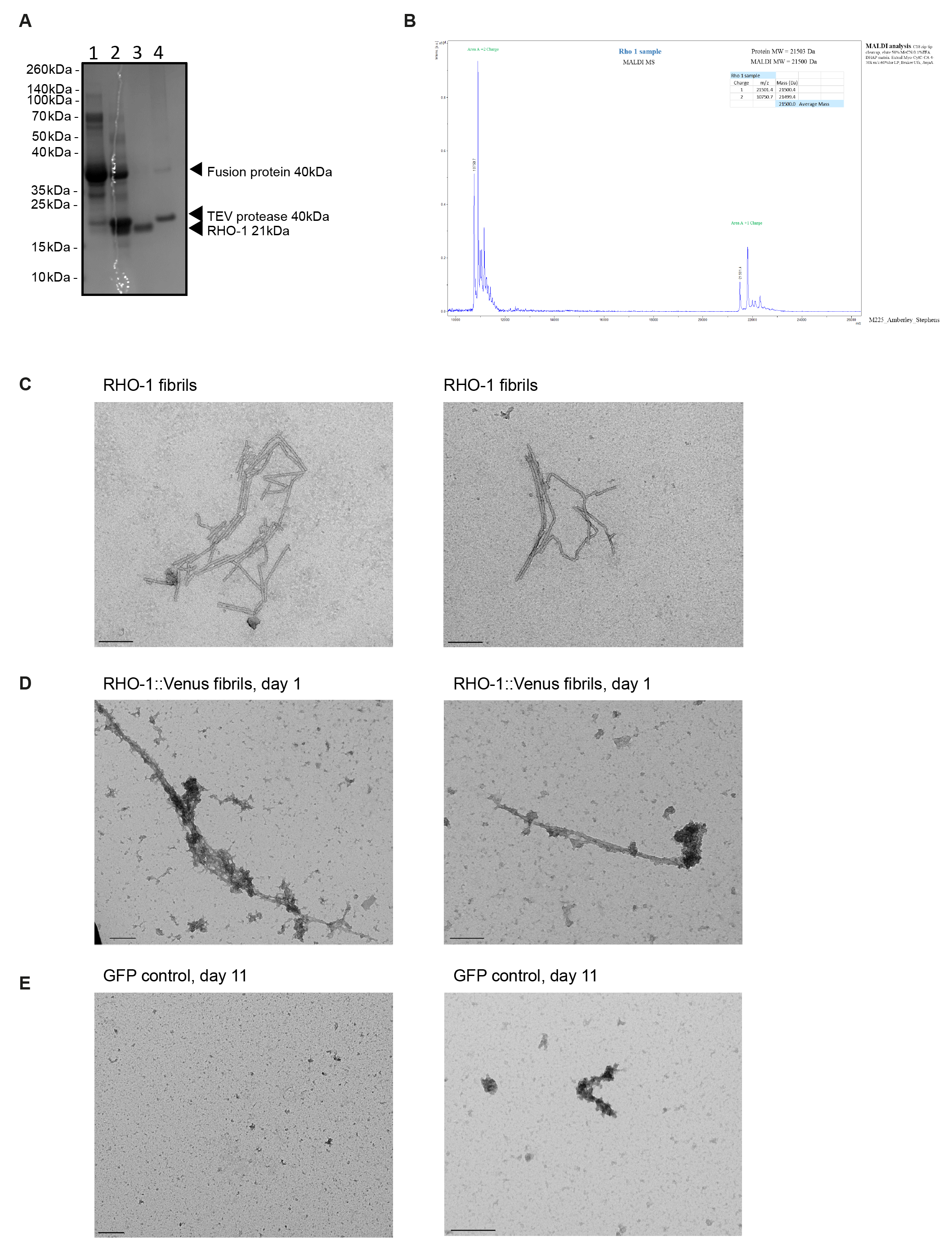
Recombinant RHO-1 forms amyloid-like fibrils *in vitro* and *in vivo*. (A) Purification steps of recombinant RHO-1 analyzed by SDS-PAGE gel electrophoresis and Coomassie staining. The RHO-1 fusion protein consisting of the thioredoxin protein, His tag and Tobacco etch virus (TEV) cleavage recognition site was purified on a Crude FF column and eluted with imidazole. It ran slightly quicker than the expected 40 kDa predicted size (lane 1). The fusion protein was incubated with recombinant TEV protease overnight in a 1:50 ratio (lane 2). The cleaved RHO-1 protein did not bind to the Crude FF column and eluted in the flow through (lane 3). The TEV protease which has a 6xHis tag and the remaining uncleaved fusion protein were eluted from the column with imidazole (lane 4). (B) Matrix Assisted Laser Desorption/Ionization (MADLI) mass spectrometry of purified RHO-1. Performed by the Proteomics Facility, Biochemistry Department, University of Cambridge. (C) Negative-stain transmission electron micrographs of fibrillised RHO-1. TEM of RHO-1 negatively stained using 2% uranyl acetate. RHO-1 was incubated for 1 week in a ThT assay to fibrillise. Scale bar = 200 nm. (D) Negative-stain transmission electron micrographs of *pmyo-2∷RHO-1∷Venus* worm lysates, day 1. TEM of RHO-1 fibrils negatively stained using 2% uranyl acetate. Scale bar = 200 nm. (E) Negative-stain transmission electron micrographs of GFP control worm lysates (CL2122), day 11. Scale bar = 200 nm.

**Supplementary figure 5:**
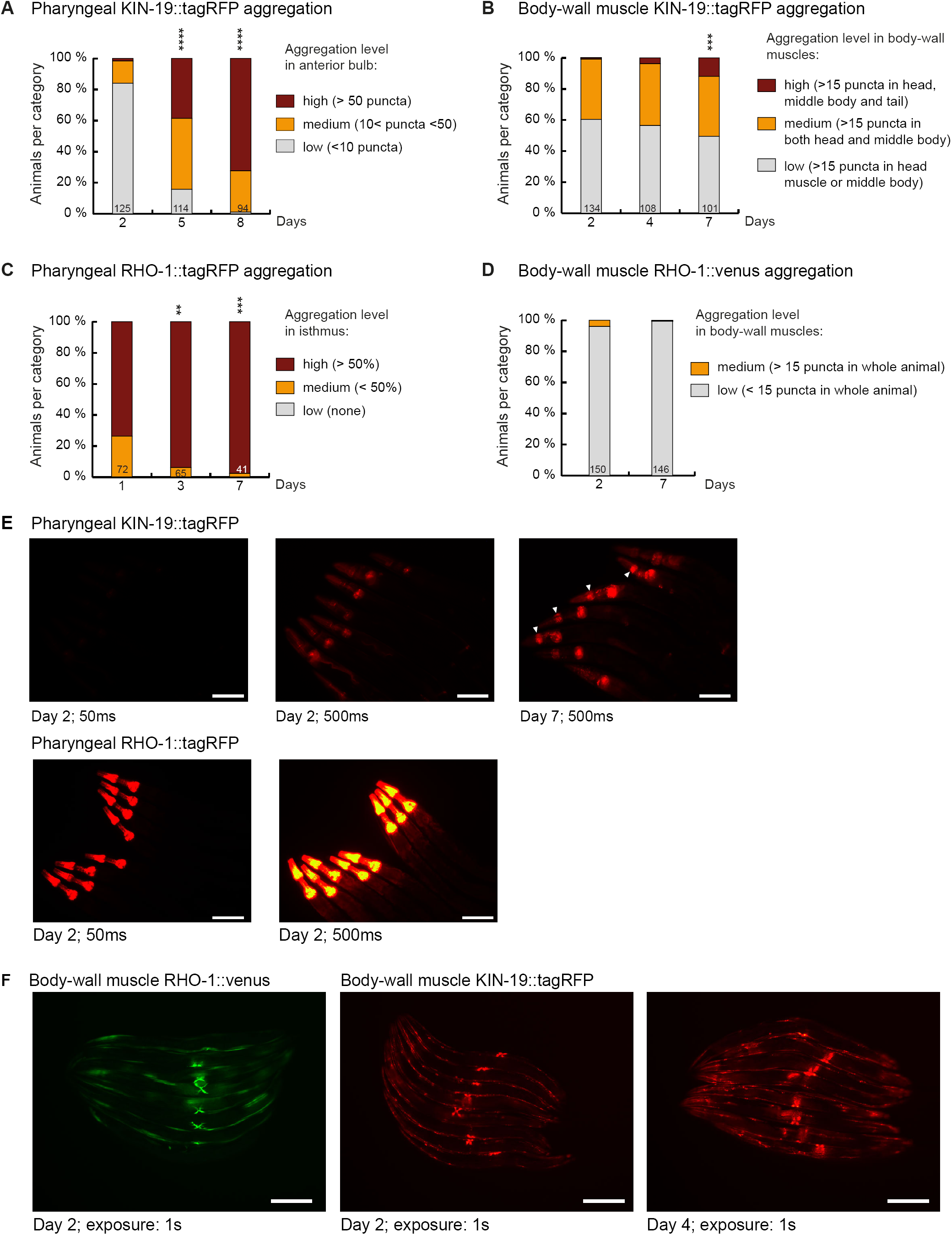
Markers for age-dependent protein aggregation. (A) Increased pharyngeal KIN-19 aggregation with age in animals expressing *Pkin-19∷KIN-19∷tagRFP*. (B) Increased body-wall muscle KIN-19 aggregation with age in animals expressing *Pmyo-3∷KIN-19∷tagRFP*. (C) Increased pharyngeal RHO-1 aggregation with age in animals expressing *Pmyo-2∷RHO-1∷tagRFP*. Already young adults display abundant RHO-1 aggregation. (D) Absence of RHO-1 aggregation in the majority of animals expressing *Punc-54∷RHO-1∷Venus*. Numbers of worms indicated in the bars. Significance calculated low + medium versus high aggregation levels compared to day 2 of adulthood; Fisher’s exact test: **p< 0.01, ***p< 0.001, ****p< 0.0001 (E) RHO-1∷tagRFP aggregates abundantly at day 2 compared to KIN-19∷tagRFP. Fluorescent micrograph of the upper body region of animals expressing *Pkin-19∷KIN-19∷tagRFP* or *Pmyo-2∷RHO-1∷tagRFP*. Days of adulthood and exposure times displayed under images. Arrow heads highlight animals with high levels of KIN-19 aggregation in anterior bulb. Scale bar: 100 μm (F) KIN-19∷tagRFP but not RHO-1∷Venus aggregates in body-wall muscle (small puncta). Fluorescent micrograph of whole animals expressing *Punc-54∷RHO-1∷Venus* or *Pmyo-3∷KIN-19∷tagRFP*. Head region on the left. Days of adulthood and exposure times displayed under images. Scale bar: 200 μm

**Supplementary Figure 6:**
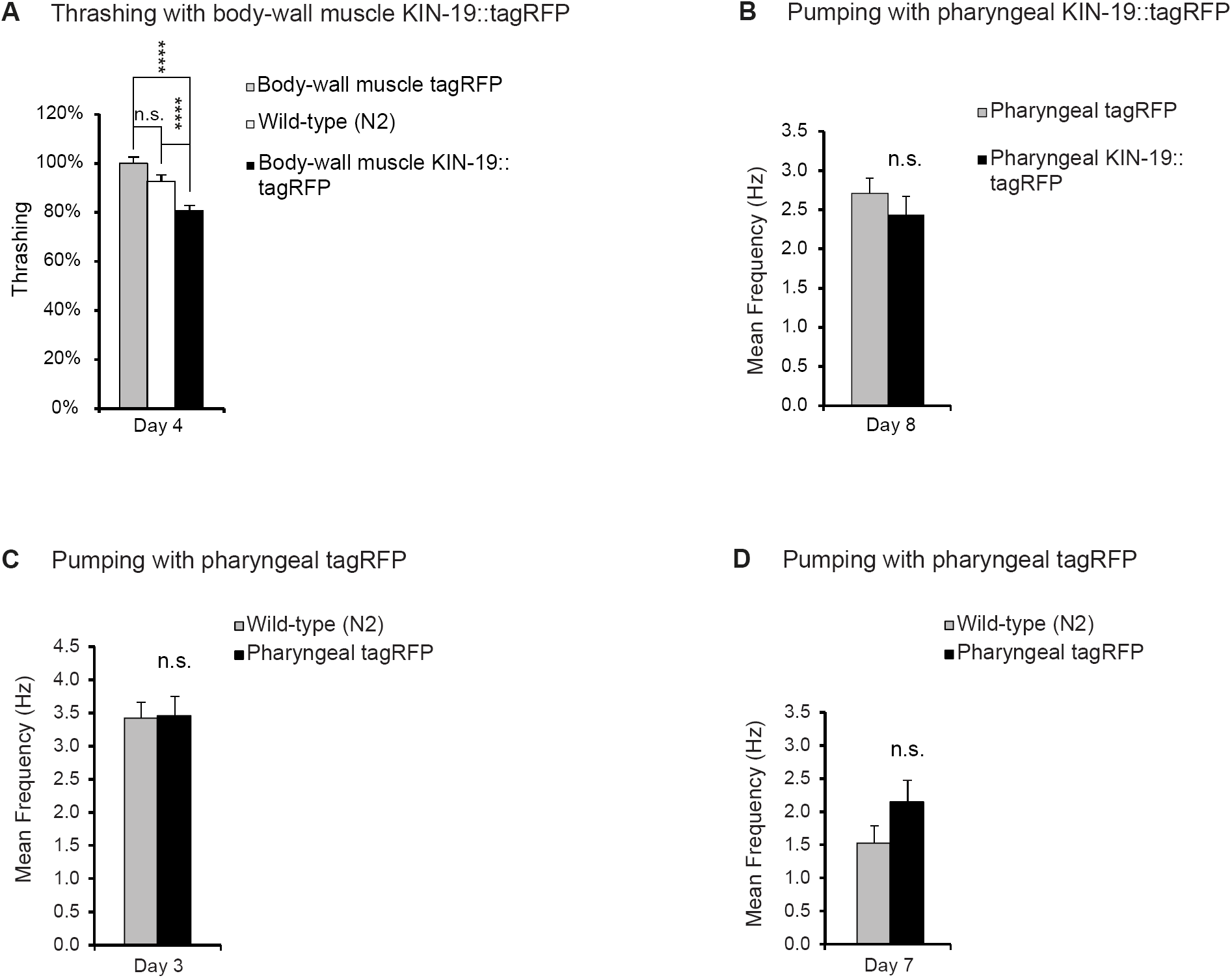
KIN-19 aggregation in body-wall muscles impairs thrashing. (A) KIN-19 aggregation in body-wall muscles impairs thrashing. Mean body bends per seconds are set to 100% in *Pmyo-3∷tagRFP* transgenics. Mann-Whitney test: ****p<0.0001 (B) No effect on pumping detected in animals with mixed levels of KIN-19∷tagRFP aggregation. N= 23-39 animals analyzed per group. T-test: non-significant. (C) Fluorescent tagRFP does not affect pharyngeal pumping in young animals. N= 23-28 animals analyzed per group. T-test: non-significant. (D) Fluorescent tagRFP does not affect pharyngeal pumping in aged animals. N= 23-28 animals analyzed per group. T-test: non-significant. SEM represented, data in Table S1.

**Table S1:**
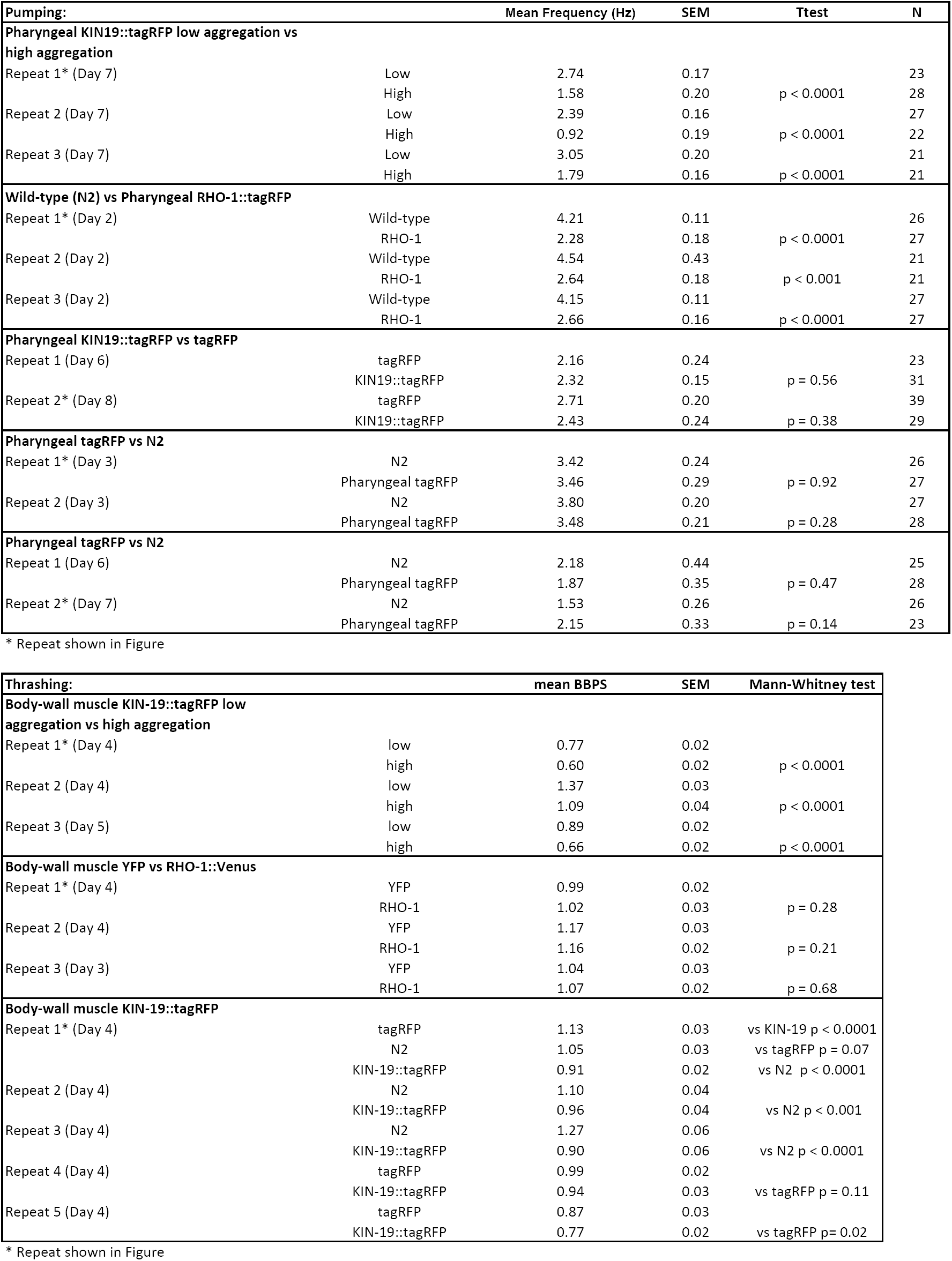
Pharyngeal pumping and thrashing repeats

## Supplementary Methods

### Pharyngeal pumping analysis

To record the electrical activity of pharynx pumping, the NemaMetrix ScreenChip System (NemaMetrix, Eugene OR) was used. The entire setup is housed in a laboratory that maintained a temperature of approximately 21°. Baseline noise was typically between 5 and 25 uV.

For each experiment, 50 worms were picked in 1.5ml of M9 + 0.01% Triton and washed 3 times via low-speed centrifugation. Worms were resuspended in 1.5ml M9 + 0.01% Triton +10uM 5-Hydroxytryptamine creatinine sulfate complex (Serotonin creatinine sulfate monohydrate) (Sigma, H7752) and incubated for 20 minutes. The ScreenChip system was placed on a stereoscope and loaded with a fresh screen chip. The screen chip was then vacuum-filled with M9 + 0.01% Triton +10uM 5-Hydroxytryptamine creatinine sulfate complex and the NemAquire software initiated for baseline noise checking. The animals were loaded into the recording channel of the screen chip via vacuum. After loading each animal, we waited at least 30 seconds or until the pumping became regular before starting to record. Each animal was recorded for approximately 2 minutes regardless of whether pumping activity was observed or not. Between 20-40 animals were recorded for each condition.

The recordings were analysed by NemAnalysis v0.2 software using the “Brute Force” optimization method. The ideal settings were chosen automatically from all combinations of the bounds settings (Minimum SNR from 1.4 (low) to 2.0 (high), with a Step size of 0.1; Highpass Cutoff from 10 (low) to 20 (high), with a Step size of 5) and applied to produce the analysis results. Data was exported into Excel for statistical analysis. The student’s t test was used for statistical analysis.

### Thrashing analysis

To quantify movement in terms of body-bends-per-second, movies of worms swimming in liquid were obtained with high frame rates (15 frames per second). A high-resolution monochrome camera (JAI BM-500 GE, Stemmer imaging GmbH, Puchheim, Germany) was used for recording movies. For each condition, between 20-60 animals were filmed. Worms were picked from cultivation plate and allowed to swim in a small plastic petri dish filled with M9 + 0.01% Triton. Petri dish containing worms was placed on a transparent platform and illuminated from bottom up with a flat backlight (CCS TH-211/200-RD, Stemmer imaging GmbH, Puchheim, Germany) to achieve homogeneous, high contrast lighting. The entire setup is housed in a laboratory that maintained a temperature of approximately 21°. Movies were taken 10 minutes after placing the animals in the liquid. Five consecutive 30-second movies were made for each group of worms. The movies were then analysed using the ImageJ wrMTrck plugin [1]. The wrMTrck plugin tracked individual worms in the movies and counted the numbers of body-bends. The input values of wrMTrck_Batch are detailed as follows:

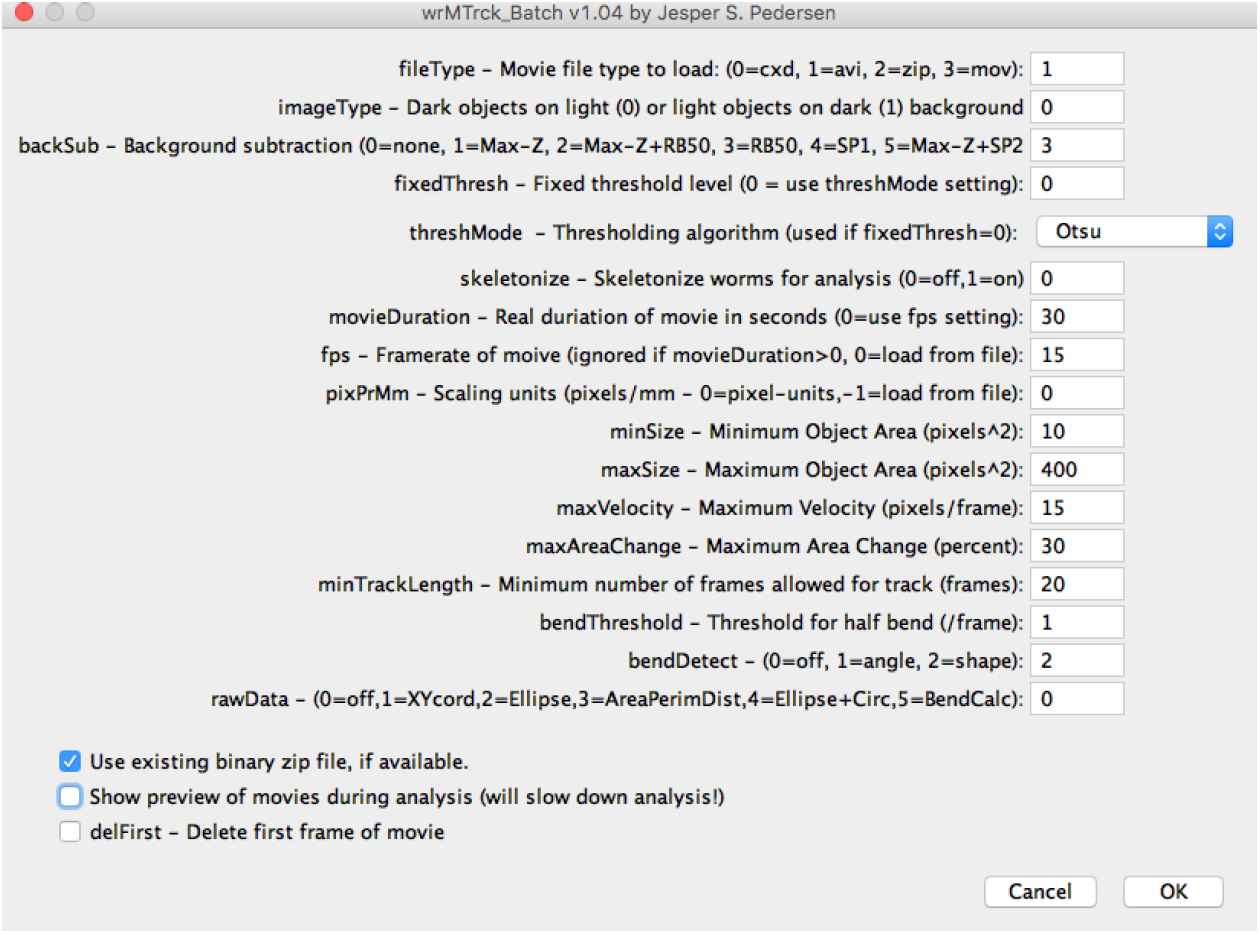

Mann-Whitney test was used for statistical analysis (GraphPad Prism 7).

### Plasmid generation for RHO-1 recombinant expression

*C. elegans* RHO-1 cDNA was cloned into pET32a expression vector using restriction sites BamHI and HindIII (NEB, UK). The open reading frame encoded the RHO-1 fusion protein comprising of a thioredoxin protein, 6xHis tag and the Tobacco etch virus (TEV) cleavage recognition site with the sequence ENLYFQA, where TEV cleaves between Q and A, which was also the N-terminal residue of the Rho-1 protein sequence, followed by the RHO-1 protein. The plasmid was confirmed by DNA sequencing (Source Bioscience, Cambridge, UK).

### Expression and Isolation of Recombinant RHO-1 from Inclusion Bodies

Reagents were purchased from Sigma-Aldrich, UK unless stated. BL21 DE3 STAR™ E. coli (Thermo Fisher Scientific, USA) were transformed with pET32a:RHO-1. 3 L cultures of E. coli in Lysogeny Broth containing carbenicillin (100 ug/mL) were grown at 37°C at 250 rpm and induced for expression of the RHO-1 fusion protein with 1 mM isopropyl-β-thiogalactopyranoside (IPTG) for four hours. *E. coli* were pelleted by centrifuge at 8000 x g for 15 minutes before being washed by resuspension in PBS with 1% Triton X-110 and 1 mM Tris(2-carboxyethyl)phosphine hydrochloride (TCEP) and centrifuged again. The pelleted *E. coli* were either stored at -20°C until further use or lysed straight away. The RHO-1 fusion protein forms in inclusion bodies. To release the inclusion bodies, the *E. coli* were resuspended in 20 mL of lysis buffer per 1 L of culture (50 mM Tris, 500 mM NaCl, 5 mM MgCl2, 1 mM phenylmethylsulfonyl fluoride (PMSF), protease inhibitor tablets (cOmplete™, Mini EDTA-free, Roche), 1% Triton X-110, 1 mM TCEP, pH 8 at 4°C) and sonicated on ice using three rounds of 30 s on and 30 s off 70% sonication power. The suspension was centrifuged at 10,000 x g for 10 minutes at 4°C. The pellet containing RHO-1 inclusion bodies was resuspended in 30 mL per 1 L culture of wash buffer 1 (Lysis buffer + 2 M urea, pH 8 at 4°C). The inclusion bodies were sonicated for four rounds of 10 s on, 20 s off and centrifuged at 10,000 x g for 10 minutes at 4°C, this wash was then repeated again. Wash number 3 and 4 used the same sonication and centrifuge parameters, but pellets were washed with wash buffer 2 (Lysis buffer, 2 M urea, 1 mM TCEP, without Triton-110 or protease inhibitors). The final pellet became paler and more chalk-like. These wash steps are important to lead to a purer final RHO-1 protein. The inclusion bodies were then solublised in solublising buffer using 10 mL per 1 L of culture for 1 hour using a magnetic stirrer (50 mM Tris, 500 mM NaCl, 5 mM MgCl2, 6 M guanidinium hydrochloride (GuHCl), 1 mM TCEP, (6 M Urea can also be used in place of GuHCl, but GuHCl gives a slightly higher final protein yield)). Insoluble material was removed by centrifuge at 16,000 x g for 10 minutes at room temperature (RT).

### Purification of Recombinant RHO-1

The RHO-1 fusion protein was purified using two linked-together 1 mL HisTrap Crude FF columns on an ÄKTA Pure (GE Healthcare, Sweden). The columns were equilibrated with solublising buffer before 10 mL of protein was loaded onto the column by the sample pump. The columns were washed in wash buffer (50 mM Tris, 500 mM NaCl, 5 mM MgCl2, 6 M Urea, 20 mM imidazole) before being eluted against a linear gradient of elution buffer (50 mM Tris, 500 mM NaCl, 5 mM MgCl2, 6 M Urea, 500 mM imidazole) over 16 column volumes of a 1 x 1 mL column, i.e. 16 mL. Multiple purification runs were performed until all RHO-1 fusion protein was purified. Fractions from all runs containing the RHO-1 fusion protein were pooled and dialysed overnight in 2 L stabilising buffer modified from ([2, 3] Healthcare 2018) (50 mM Tris, 250 mM NaCl, 100 mM arginine, 5 mM reduced glutathione, 0.5 mM oxidised glutathione, 5 mM MgCl2, 5 μM guanosine 5’-diphosphate (GDP) (Generon, UK) pH 7.2). The RHO-1 fusion protein was then concentrated to 1 mL per 1 L culture using Spectra/Gel Absorbent (Spectrum Labs, USA) through a 10 kDa MWCO Slide-A-Lyzer™ dialysis cassette (Thermo Fisher Scientific). The RHO-1 fusion protein was then incubated overnight with recombinant TEV protease to cleave the RHO-1 from the fusion tag in a 1:50 ration of TEV to RHO-1 fusion protein based on absorbance at 280 nm. Recombinant TEV was produced by Dr Marielle Wälti using methods from [4]. The cleaved RHO-1 was separated from the fusion tag and TEV protein by purification using the two linked-together 1 mL HisTrap Crude FF columns on an ÄKTA Pure. The columns were equilibrated in stabilising buffer and 1 mL of the protein loaded onto the columns by injection. Cleaved RHO-1 was eluted in stabilisation buffer during the column wash. The fusion tag and the His-tagged TEV were retained on the column and eluted with 100% elution buffer over 10 CV (stabilisation buffer with 500 mM imidazole). Protein concentration was monitored throughout purification using the extinction coefficient 0.862 M^-1^ cm^-1^ for the fusion protein, 0.849 M^-1^ cm^-1^ for the refolded fusion protein and 0.875 M^-1^ cm^-1^ for the cleaved refolded RHO-1 protein. Quantitative analysis of protein purity was performed in FIJI image analysis software [5] by profiling protein band intensity of the stained gel mass spectrometry confirmed the purification of RHO-1. To note, RHO-1 precipitated in NaP, Tris, NaCl buffers. RHO-1 protein mass was confirmed by mass spectrometry.

### Fibrillisation of RHO-1 and analysis by Transmission Electron Microscopy (TEM)

RHO-1 was fibrillised by incubating 20 μM in stabilising buffer during a Thioflavin-T (ThT) based-assay with 10 μM ThT (AbCam, UK) in non-binding, clear bottom, black 96-well plate (PN 655906 Greiner Bio-One GmbH, Germany). The plate was incubated at 37°C with constant shaking at 300 rpm for 7 days. RHO-1 was taken from the wells in the microplate for imaging by TEM. All samples also contained 0.05% NaN3 to prevent bacterial growth. Samples were incubated in an oven rotating at maximum speed (UVP HB-1000 Hybridizer, Fisher Scientific) at 37°C for five weeks.

Fibrillised RHO-1 samples were centrifuged for 20 minutes at 21,000 xg and the supernatant removed leaving 10 μL of sample. Each 10 μL sample was incubated on a glow-discharged copper grid for 1 minute. Excess liquid was blotted off and the grid washed in twice in dH2O for 15 seconds. 2% uranyl acetate was used to negatively stain the samples for 30 s before imaging on the Tecnai G2 80-200kv TEM at the Cambridge Advanced Imaging Centre.

### Preparation and analysis of worm lysates for fibrils by TEM

*C. elegans* DCD146 expressing RHO-1∷Venus were grown to confluency on high growth medium plates and bleached to obtain a synchronized population of worms as previously described [6]. L1s were transferred into a liquid culture with complete S basal supplemented with OP50-1 (OP50 with Streptomycin resistance) and grown at 20°C as previously described [7, 8]. At day 1 of adulthood, worms were allowed to sediment in a separation funnel and washed with cold M9. Worm pellet was resuspended in PBS with 2x protease inhibitor tablets (cOmplete™, Mini EDTA-free, Roche) and drip-frozen into liquid nitrogen before being ground in a mortar. Frozen ground worms were resuspended in radioimmunoprecipitation assay (RIPA) buffer with protease inhibitor tablets (cOmplete™, Mini EDTA-free, Roche) and lysed by 20-25 passages through a cell homogeniser (Isobiotec, Germany) using a tungsten carbide ball with 16 μm clearance. Cuticle fragments and unlysed worms were removed by centrifugation for 5 min at 835 g and 4 °C. After careful removal of the supernatant, the insoluble fraction was collected by centrifugation for 30 min at 21,000 g and 4 °C. The supernatant was removed and the pellet was resuspended in phosphate buffered saline (PBS) with protease inhibitor tablets (cOmplete™, Mini EDTA-free, Roche) by homogenising with a needle (27G, Sterican). 10 μL sample was applied to a carbon coated grid, and 2% uranyl acetate was used for negative staining. Imaging was performed on the Tecnai G2 80-200kv TEM at the Cambridge Advanced Imaging Centre.

Healthcare, Ge. 2018. Purifying Challenging Proteins Principles and Methods. Accessed May 15. https://cdn.gelifesciences.com/dmm3bwsv3/AssetStream.aspx?mediaformatid=10061&destinationid=10016&assetid=19487.

